# A common phthalate replacement disrupts ovarian function in young adult mice

**DOI:** 10.1101/2024.08.14.607936

**Authors:** Courtney Potts, Allison Harbolic, Maire Murphy, Michelle Jojy, Christine Hanna, Maira Nadeem, Hanin Alahmadi, Stephanie Martinez, Genoa R. Warner

## Abstract

Di-2-ethylhexyl terephthalate (DEHTP) is a replacement for its structural isomer di-2-ethylhexyl phthalate (DEHP), a known endocrine disrupting chemical and ovarian toxicant. DEHTP is used as a plasticizer in polyvinyl chloride products and its metabolites are increasingly found in biomonitoring studies at levels similar to phthalates. However, little is known about the effects of DEHTP on the ovary. In this research, we tested the hypothesis that DEHTP is an ovarian toxicant and likely endocrine disrupting chemical like its isomer DEHP. The impact of environmentally relevant exposure to DEHTP and/or its metabolite mono-2-ethylhexyl terephthalate (MEHTP) on the mouse ovary was investigated *in vivo* and *in vitro.* For the *in vivo* studies, young adult CD-1 mice were orally dosed with vehicle, 10 µg/kg, 100 µg/kg, or 100 mg/kg of DEHTP for 10 days. For the *in vitro* studies, isolated untreated ovarian follicles were exposed to vehicle, 0.1, 1, 10, or 100 µg/mL of DEHTP or MEHTP. Follicle counts, hormone levels, and gene expression of steroidogenic enzymes, cell cycle regulators, and apoptosis factors were analyzed. *In vivo*, DEHTP exposure increased primordial follicle counts at 100 µg/kg and 100 mg/kg and decreased primary follicle counts at 100 mg/kg compared to control. DEHTP exposure also decreased expression of cell cycle regulators and apoptotic factors compared to control. *In vitro*, follicle growth was reduced by 1 µg/mL DEHTP and 1, 10, and 100 µg/mL MEHTP compared to controls, and expression of the cell cycle regulator *Cdkn2b* was increased. Steroid hormone levels and steroidogenic enzyme gene expression trended toward decreases *in vivo*, whereas progesterone was significantly increased by exposure to 100 µg/mL MEHTP *in vitro*. Overall, these results suggest that DEHTP and MEHTP may be ovarian toxicants at low doses and should be subjected to further scrutiny for reproductive toxicity due to their similar structures to phthalates.

## 1. Introduction

Plastics are some of the most versatile and widely used materials in the world, greatly improving the quality of life for humans. Plasticizers, which make plastics more flexible and improve durability, are an essential additive. However, an extensive body of research now exists to show that ortho-phthalate plasticizers, such as di-2-ethylhexyl phthalate (DEHP), are reproductive toxicants and endocrine disrupting chemicals (EDCs) [1–4]. In response to restrictions on its use in products and consumer pressure, manufacturers have moved to replace DEHP with alternative chemicals such as di-2-ethylhexyl terephthalate (DEHTP), a para-substituted structural isomer (**Figure 1**) [5]. Its primary hydrolysis metabolite is mono-2- ethylhexyl terephthalate (MEHTP), an isomer of the bioactive DEHP metabolite mono-2- ethylhexyl phthalate (MEHP).

**Figure 1:**
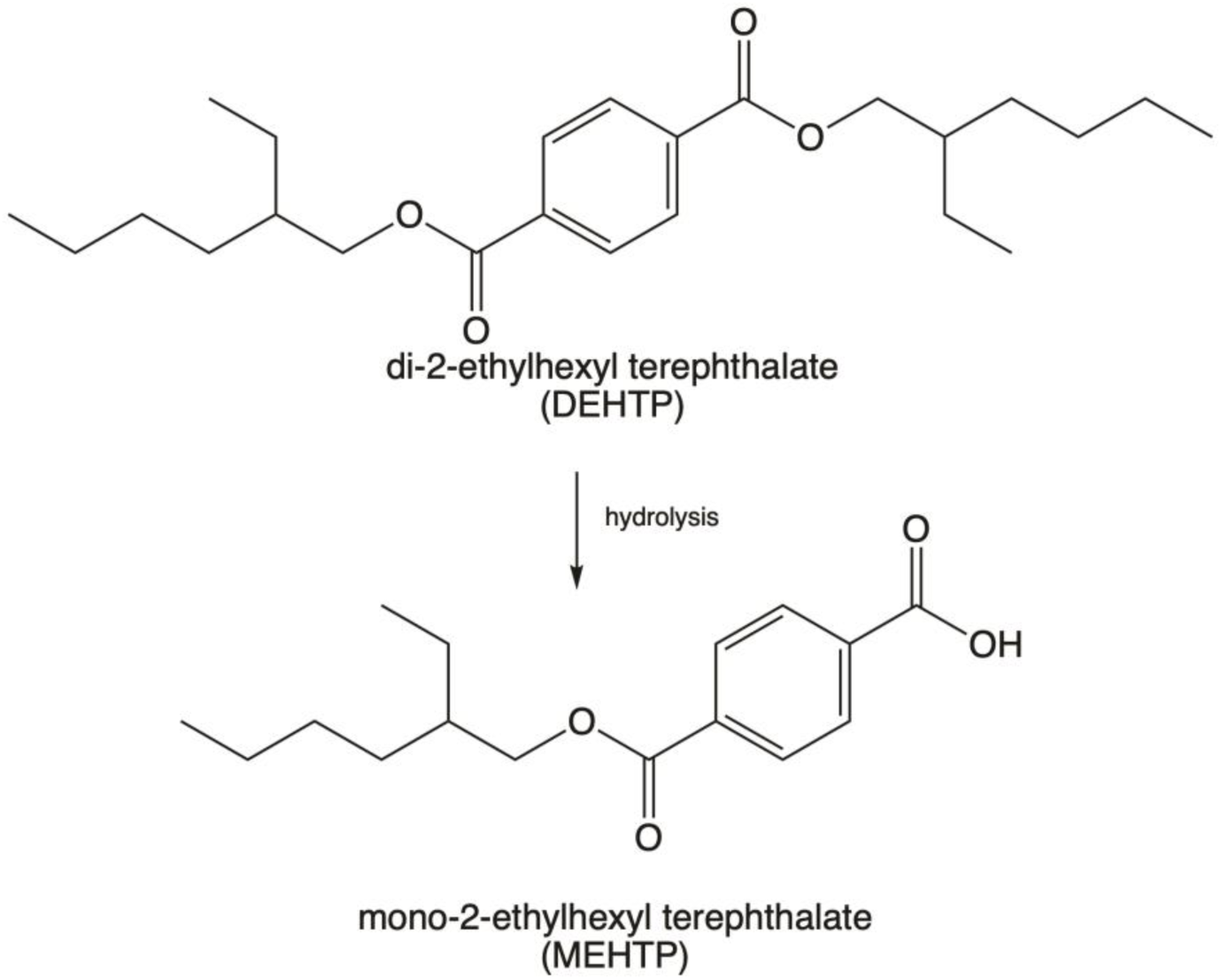
Structure of the para-substituted ester di-2-ethylhexyl terephthalate (DEHTP) and its primary metabolite mono-2-ethylhexyl terephthalate (MEHTP).

Since the introduction of phthalate alternatives in the early 2000s, DEHTP biomarker concentrations have increased to levels comparable to DEHP metabolites in US adults [6]. From 2009 to 2016, adults in the US showed an average 51% increase per year in the urinary levels of DEHTP metabolites [7]. DEHTP metabolites have been detected in children and pregnant women around the world [8–13] and analysis of global populations confirm widespread and increasing exposure of DEHTP on par with DEHP and other phthalates [7,14]. It has been detected at high levels in fast food, infant formula, and children’s toys, and biomonitoring studies suggests that it may be used in cosmetic or fragranced products [15–20]. Of particular concern are reports of high concentrations of DEHTP metabolites in adult women, including during pregnancy, due to occupational exposure and/or greater use of personal care products compared to men [8,15,21]. A recent study of pregnant women in Illinois found higher concentrations of DEHTP metabolites than any other measured phthalate or phthalate alternative, emphasizing the broad market share of DEHTP and the pressing need to understand its reproductive toxicity potential [21].

Epidemiology studies suggest that DEHTP exposure may disrupt hormone levels and cause other reproductive alterations in humans. In women samples through the US Center for Disease Control’s National Health and Nutrition Examination Survey (NHANES), higher concentrations of DEHTP metabolites were associated with increased testosterone-to-estradiol ratio in post- menopausal women [22]. In the Illinois study of pregnant women, DEHTP metabolites were associated with increased maternal estrogen levels and caused sex-specific effects, with higher estrogen in late gestation in women carrying female fetuses and lower estrogen in early gestation in women carrying male fetuses, similar to the observed associations with DEHP metabolites [23]. In another cohort of pregnant women in the US, DEHTP metabolites were associated with oxidative stress and inflammation [24].

Very few studies have investigated the reproductive toxicity of environmentally relevant doses of DEHTP in *in vitro* or animal models. A human H295R cell line steroidogenesis assay found that estradiol and testosterone production were disrupted by DEHTP’s monoester metabolites [25]. Female zebrafish exposed to DEHTP for 21 days laid fewer eggs than controls and both sexes had altered levels of estradiol and/or testosterone [26]. In this study, we use *in vitro* and *in vivo* rodent models to assess the ovarian toxicity of DEHTP and its primary metabolite MEHTP. The ovary is highly conserved in mammals, thus providing translatability to human health [27]. This study focuses on the primary ovarian functions of steroid hormone production (steroidogenesis) and follicle development (folliculogenesis) to provide insight into the potential of DEHTP to act as an ovarian toxicant at human-relevant doses.

## 2. Materials and methods

### 2.1 Chemicals

DEHTP was obtained from Sigma Aldrich (catalog no. 525189, ≥ 96% purity) and MEHTP from LGC Standards (TRC-M542570, 98% purity). For the *in vivo* study, mice were dosed with vehicle (corn oil) or 10 µg/kg, 100 µg/kg, or 100 mg/kg of DEHTP for 10 days. For the *in vitro* studies, follicles were exposed to vehicle control (dimethyl sulfoxide, DMSO, Sigma Aldrich D2650, ≥ 99.7% purity) or 0.1, 1, 10, or 100 µg/mL of DEHTP or MEHTP. The doses were chosen for environmental relevance and for comparison to previous studies with phthalates in the ovary [3]. Metabolites of DEHTP in human urine are comparable to phthalate metabolite levels [21,28]. The 10 µg/kg dose and the 0.1 and 1 µg/mL concentrations are representative of everyday human exposure; the 100 µg/kg dose and 10 µg/mL concentration are representative of occupational or high use exposure; the 100 mg/kg dose and 100 µg/mL concentrations are high doses for comparison to previous studies with phthalates and for providing mechanistic insight [29]. Treatment solutions of 0.13, 1.33, 13.3, and 133.3 mg/mL of DEHTP and MEHTP were prepared by diluting each chemical in DMSO such that the same volume of the treatment solution (0.75 µL) could be added per mL of culture media to maintain consistent DMSO concentration of 0.075%.

### 2.2 Animals and in vivo study design

Young adult female CD-1 mice were purchased from Charles River (Wilmington, Massachusetts) at 23 days of age. Mice were housed in the College of Veterinary Medicine Animal Facility at the University of Illinois at Urbana-Champaign and allowed to acclimate to the facility prior to experimentation. Temperature was maintained at 22 ± 1 °C with 12 h light-dark cycles to provide a controlled housing environment. Food and water were provided ad libitum. At 30 days of age, mice were orally dosed by pipetting the solution into the mouth with either vehicle control (corn oil) or DEHTP (10 µg/kg, 100 µg/kg, or 100 mg/kg) for 10 days. Dosing volumes were determined daily by corresponding mouse weight. Mice were euthanized via carbon dioxide inhalation during diestrus for sample collection within 5 days of the final dose. Cyclicity was assessed daily at the same time via vaginal lavage. Body weight and ovary, uterus, and liver weights were determined following euthanasia. Blood was collected from the inferior vena cava after euthanization, allowed to sit at room temperature for 15 min, and then centrifuged to isolate sera. Sera were frozen at -80 °C until analysis of sex steroid hormone levels. Ovaries were collected for follicle counts and gene expression analyses (1 ovary each, randomly assigned). All procedures involving animal care, euthanasia, and tissue collection were approved by the Institutional Animal Use and Care Committee at the University of Illinois at Urbana-Champaign.

### 2.3 Follicle counting

Following exposure to DEHTP for 10 days, ovaries were fixed in Dietrich’s fixative, transferred to 70% ethanol, dehydrated, embedded in paraffin wax, and serially sectioned (8 µm) using a microtome. Sections were stained with eosin and hematoxylin and every 10th section was counted. Stages of follicular development were analyzed following previous criteria [29–31]. In brief, primordial follicles contain the oocyte and are surrounded by a single layer of granulosa cells. Primary follicles contain the oocyte while surrounded by cuboidal layer of granulosa cells. Preantral follicles have 2 or more layers of cuboidal granulosa cells and an outer layer of theca cells. Finally, antral follicles have multiple layers of theca and granulosa cells and a fluid filled space within the follicle. Follicles were considered abnormal if the oocyte appeared shriveled and darkly stained. Primordial and primary follicles were all counted regardless of nuclear material within the oocyte, whereas preantral and antral follicles were only counted when the nuclei were visible to avoid duplicate counting. Follicles that appear to be transitioning between stages were counted as the more immature phase. Follicle counting was performed blinded to treatment group.

### 2.4 Antral follicle culture

Untreated adult female CD-1 mice were euthanized at 32–38 days of age. Antral follicles were manually isolated from the ovaries based on relative follicle size (200–400 µm). Interstitial tissue was cleaned using watchmaker forceps. Follicles were obtained from 3-4 mice and were individually plated in 96 well culture plates with 10-12 follicles per group. The different treatment groups (DMSO or 0.1–100 µg/mL) were supplemented in minimum essential medium alpha (Gibco) containing 1% insulin-transferrin-selenium (10 ng/ml insulin, 5.5 mg/ml transferrin, 5 ng/ml sodium selenite, Sigma-Aldrich), 100 U/ml penicillin (Sigma-Aldrich), 100 mg/ml streptomycin (Sigma-Aldrich), 5 IU/mL human recombinant follicle-stimulating hormone (Dr A.F. Parlow, National Hormone and Peptide Program, Harbor-UCLA Medical Center, Torrance, California), and 5% fetal bovine serum (Atlanta Biologicals, Lawrenceville, Georgia). Stock concentrations of DEHTP and MEHTP (0.1, 1, 10, 100 µg/mL) were prepared, and 0.75 µL of stock was used per 1 mL of supplemented medium. Follicles were cultured for 96 h or 24 hr in an incubator supplying 5% CO2 at 37 °C.

### 2.5 Follicle growth analysis

Follicle growth was assessed every 24 hours for the 96 hour culture period by measuring follicle diameters on perpendicular axes with an inverted microscope equipped with a calibrated ocular micrometer. The diameters of each follicle were averaged within the treatment group for each 24 hr interval, and the average values were divided by the initial average measurement at 0 hr of each of the respective treatment groups to calculate the percent change in follicle diameter over time. This percent change in antral follicle diameter over time was used for statistical analysis.

### 2.6 Hormone assays

After the 96 and 24 hr follicle cultures were completed, media were pooled by treatment group and frozen at -80 °C until further analysis. Hormone analysis was performed on media from in vitro experiments and serum from in vivo experiments via enzyme linked immunosorbent assay (ELISA). Kits for estradiol, testosterone, progesterone, androstenedione, and pregnenolone were purchased from DRG International, Inc. (Mountainside, New Jersey) and used according to the manufacturer’s instructions. Samples were run in duplicate. Samples were diluted as needed to match the dynamic range of each ELISA kit and read with a Tecan Spark multimode microplate reader or an Agilent Biotek Epoch microplate reader. The detection ranges were 0.0083–16 ng/ml for testosterone, 0–40 ng/ml for progesterone, 10.6–2000 pg/ml for estradiol, and 0–10 ng/ml for androstenedione. Hormone concentrations were interpolated using Graphpad Prism software. The mean values of each sample were used for statistical analysis.

### 2.7 Gene expression analysis

After the 96 and 24 hr follicle cultures were completed, follicles were pooled by treatment group and frozen at -80 °C until further analysis. Total RNA was extracted from the follicle using an E.Z.N.A. MicroElute®Total RNA Kit (Omega Bio-tek, Norcross, Georgia) following manufacturer’s instructions. The concentrations of RNA collected were measured using a Nanodrop spectrophotometer. cDNA was synthesized from 100 ng RNA using an iScript RT kit (Bio-Rad Laboratories, Inc., Hercules, California). The qPCR reactions were run in duplicate with 1.67 ng cDNA and 7.5 pmol gene-specific primers (Integrated DNA Technologies, Inc. Coralville, Iowa) for a final reaction volume of 10 µl. Expression levels of aromatase (*Cyp19a1*), 17a- hydroxylase (*Cyp17a1*), 17b-hydroxysteroid dehydrogenase 1 (*Hsd17b1*), cytochrome P450 11A1 (*Cyp11a1*), 3-beta-hydroxysteroid dehydrogenase 1 (*Hsd3b1*), steroidogenic acute regulatory protein (*Star*), Beta actin (*ActB),* caspase 3 (*Casp3)*, caspase 8 (*Casp8*), Bcl2-like 10 *(Bcl2l10*), B cell leukemia/lymphoma 2 *(Bcl2*), BH3 interacting domain death agonist (*Bid*), Bcl2-associated X protein (*Bax*), Bcl2-associated agonist of cell death (*Bad*), antigen identified by monoclonal antibody Ki 67 (*Ki67),* cyclin dependent kinase inhibitor 2A (*Cdkn2a*), cyclin dependent kinase inhibitor 1A (*Cdkn1a*), cyclin dependent kinase inhibitor 2B (*Cdkn2b)*, cyclin A2 (*Ccna2*), cyclin dependent kinase 4 (*Cdk4*), cyclin dependent kinase 2 (*Cdk2*), cyclin dependent kinase inhibitor 1B (*Cdkn1b*), cyclin D2 (*Ccnd2*), cyclin D1 (*Ccnd1*), and 18S ribosomal RNA (*Rn18s*) were analyzed and compared among different treatment groups (**Table 1S**). *Rn18s* was used for normalization of *in vivo* samples [32,33], and *ActB* was used as a reference gene for normalization for *in vitro* samples [34] because *Rn18s* was significantly altered by *in vitro* DEHTP exposure (**Figure S1**). *Rn18s* was retained as a housekeeper *in vivo* because it had less variability by treatment group than *ActB*. Following the completion of qPCR, the Pfaffl method was used to obtain relative fold changes in comparison to controls [35].

### 2.8 Statistical analysis

Data were expressed as the mean ± standard error of the mean (SEM). Missing values from undetectable samples were imputed as the minimum detectable value divided by the square root of 2 for ELISAs and as the maximum detectable cycle value (36) for qPCR. Data were analyzed by comparing treatment groups to control using IBM SPSS version 29 software (SPSS Inc., Chicago, IL, USA). All data were continuous and assessed for normal distribution by Shapiro-Wilk analysis. If data met assumptions of normal distribution and homogeneity of variance, data were analyzed by one-way analysis of variance (ANOVA) followed by Tukey HSD or Dunnett 2- sided *post-hoc* comparisons. However, if data met assumptions of normal distributions, but not homogeneity of variance, data were analyzed by ANOVA followed by Games-Howell or Dunnett’s T3 *post-hoc* comparisons. If data were not normally distributed or presented as percentages, the independent sample Kruskal-Wallis H followed by Mann-Whitney U non-parametric tests were performed. For all comparisons, statistical significance was determined by p-value ≤ 0.05. If p-values were greater than 0.05, but less than 0.10, data were considered to exhibit a trend towards significance.

## 3. Results

### 3.1 Effects of DEHTP exposure on body weight and organ weight in vivo

Oral exposure to DEHTP for 10 days statistically significantly increased liver weight normalized to body weight in the 10 µg/kg treatment group only (**Figure S2**). Exposure to DEHTP did not alter raw (unnormalized) ovary, uteri, liver, or body weight at collection compared to control. Normalized ovary and uteri weights were also not significantly different from control. Initial body weights were statistically indistinguishable between groups and body weight changes over the experiment were not statistically significant between groups (data not shown).

### 3.2 Effects of DEHTP exposure on follicle counts and percentages in vivo

DEHTP exposure at 100 mg/kg significantly decreased raw counts of primordial follicles and significantly increased raw counts of primary and abnormal follicles compared to vehicle controls (**Figure 2A**). When follicle counts were analyzed as percentages of the total follicle population, there was a significant decrease in primordial follicles at 100 mg/kg compared to control. Primary follicles increased at 100 µg/kg and 100 mg/kg compared to control. Preantral follicles borderline statistically significantly decreased at 100 µg/kg compared to control, and abnormal follicles significantly increased at 100 mg/kg compared to control.

**Figure 2:**
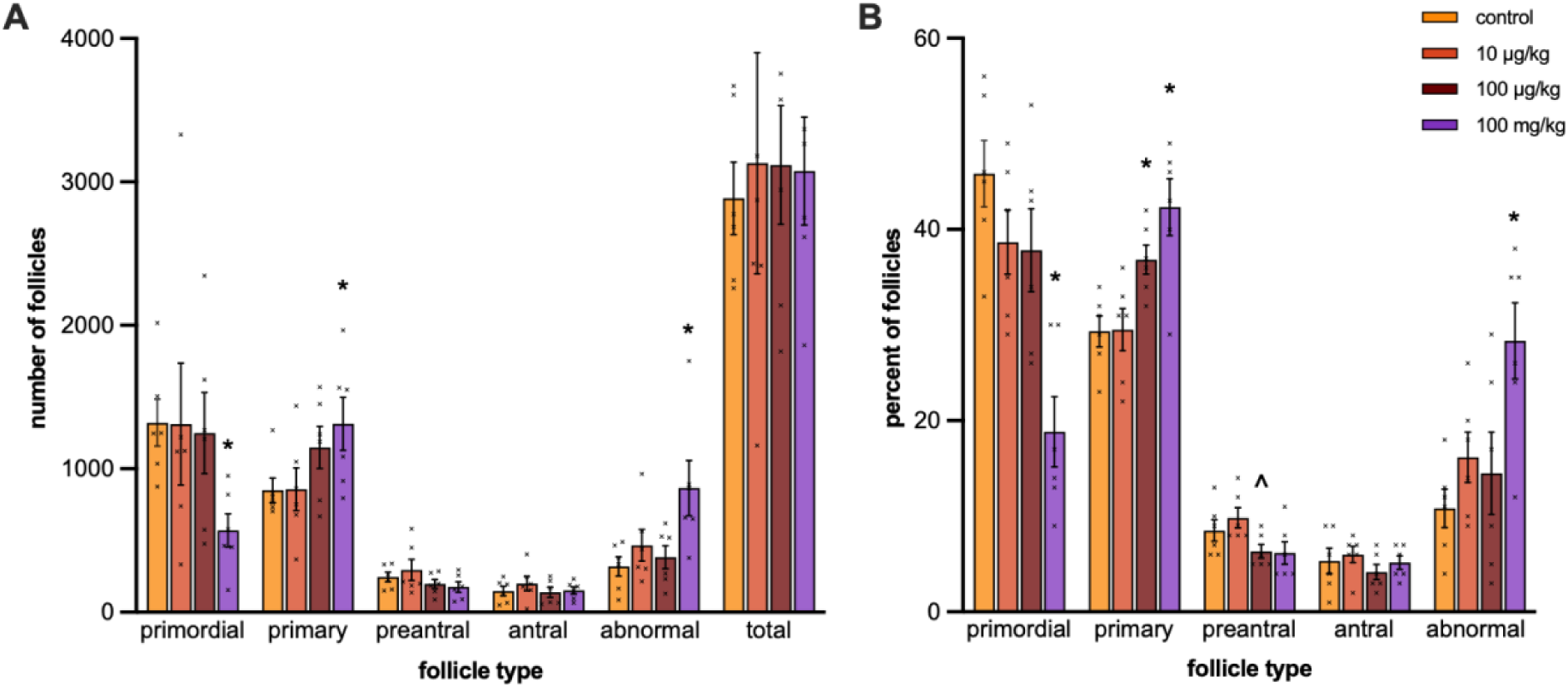
Effect of adult exposure to DEHTP on total follicle numbers (A) and percentage of each follicle type (B) in mice. Ovaries were subjected to histological examination of follicle numbers. Graphs represent means ± SEM from 5-6 animals per treatment group. Asterisks (∗) indicate significant differences from the control (p ≤ 0.05) and ^ indicates a trend toward significance (p ≤ 0.10).

### 3.3 Effects of DEHTP exposure on cell cycle and apoptosis regulators in vivo

Exposure to 100 µg/kg DEHTP borderline statistically significantly decreased expression of cell cycle regulators *Ki67* and *Ccnb1* compared to control (**Figure 3A**). Exposure to 100 mg/kg DEHTP statistically significantly decreased expression of *Ccnb1, Ccnd2, Cdk2*, and *Cdk4* compared to control. Exposure to 100 mg/kg DEHTP also statistically significantly decreased expression of apoptosis regulators *Bax* and *Bad* and borderline decreased *Bcl2* and *Bid* compared to control (**Figure 3B**).

**Figure 3:**
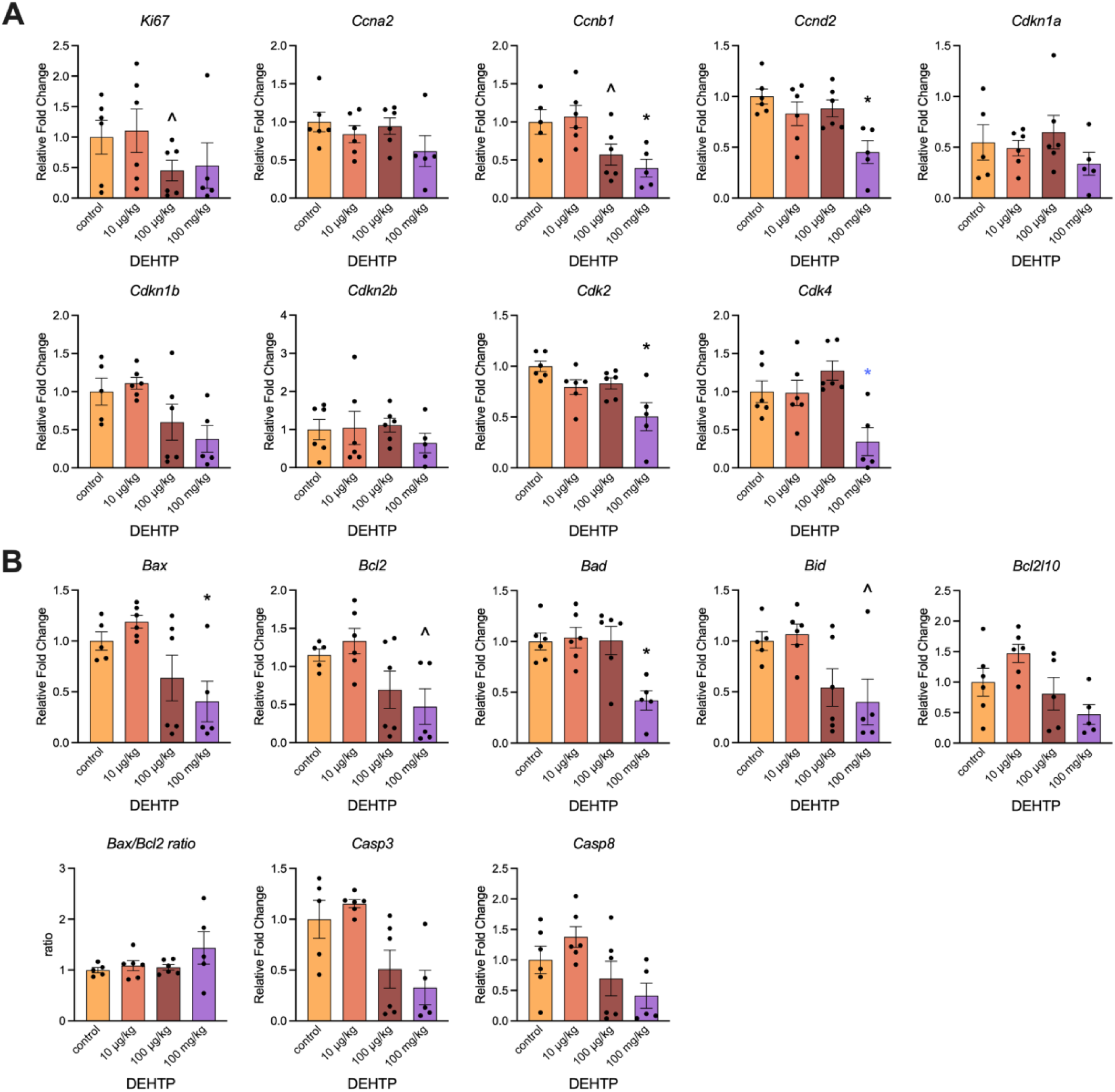
Effects of *in vivo* DEHTP exposure on cell cycle (A) and apoptosis regulator (B) gene expression in ovaries. Relative fold changes of each gene normalized to *Rn18s* are shown. Ratio of the gene expression was calculated and is shown for *Bax/Bcl2*. Graphs represent means ± SEM from 5-6 animals per treatment group. Asterisks (∗) indicate significant differences from the control (p ≤ 0.05) and ^ indicates a trend toward significance (p ≤ 0.10).

### 3.4 Effects of DEHTP exposure on sex steroid hormone levels and steroidogenic enzyme gene expression in vivo

Exposure to 100 µg/kg and 100 mg/kg DEHTP borderline statistically significantly decreased serum testosterone levels compared to control (**Figure 4A**). Only testosterone and progesterone were measured due to low serum volumes. Gene expression analysis of steroidogenic enzymes identified borderline decreases in *Cyp11a1* at 100 mg/kg and *Cyp17a1* at 100 µg/kg (**Figure 4B**).

**Figure 4:**
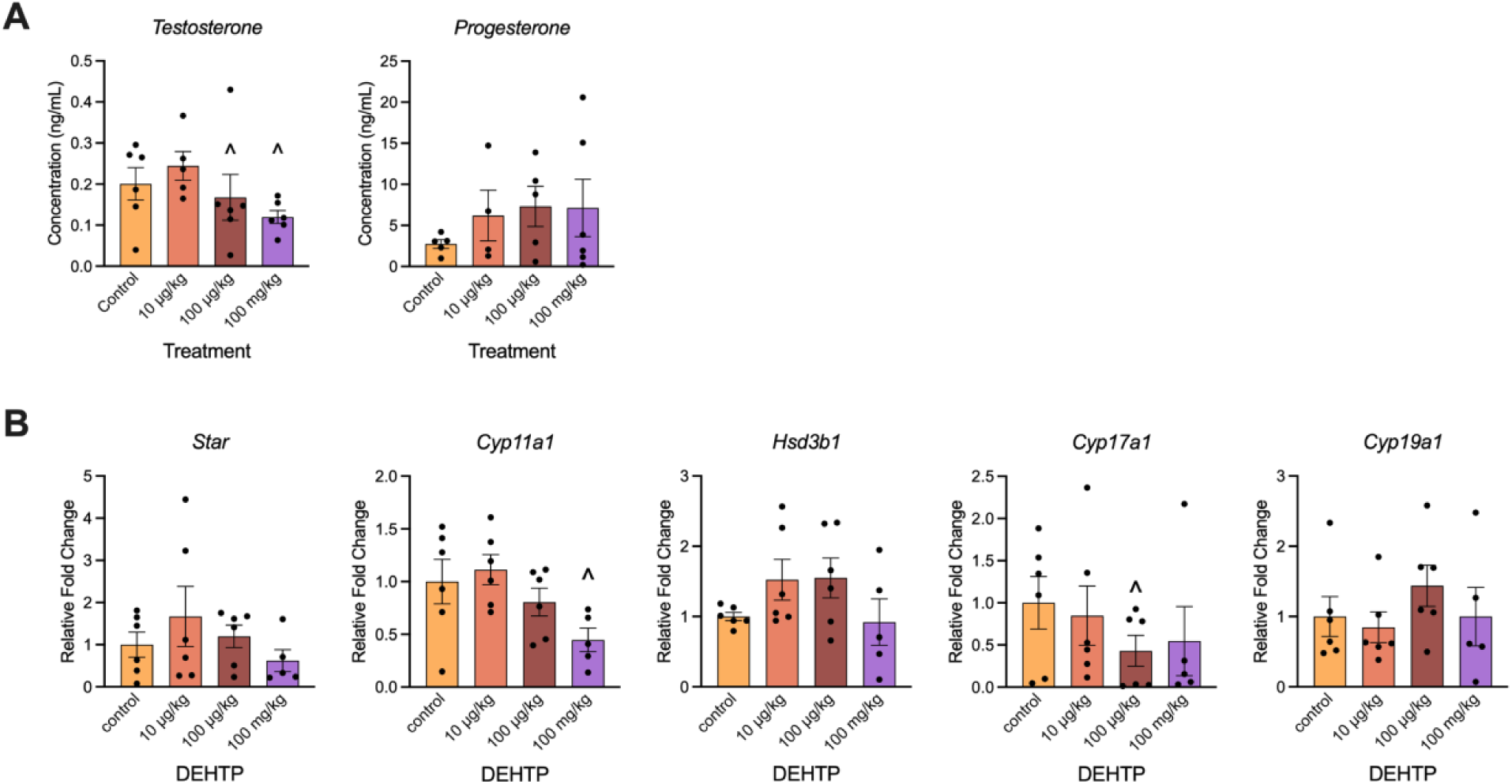
Effects of *in vivo* DEHTP exposure on serum sex steroid hormone levels (A) and steroidogenic enzyme gene expression (B). Relative fold changes of each gene normalized to *Rn18s* are shown. Graphs represent means ± SEM from 5-6 animals per treatment group. Asterisks (∗) indicate significant differences from the control (p ≤ 0.05) and ^ indicates a trend toward significance (p ≤ 0.10).

### 3.5 Effects of DEHTP and MEHTP on follicle growth in vitro

To assess the effects of DEHTP and MEHTP on follicle growth, isolated antral follicles were cultured in media containing vehicle control (DMSO) or various doses of DEHTP or MEHTP for 96 and 24 hr. DEHTP treatment at 1 µg/mL significantly decreased follicle growth at 72 and 96 hours compared to control (**Figure 5A**). MEHTP treatment also reduced follicle growth after 48 hours at 1 µg/mL, 72 and 96 hours for 10 µg/mL, and 48, 72, and 96 hours for 100 µg/mL compared to control (**Figure 5B**).

**Figure 5:**
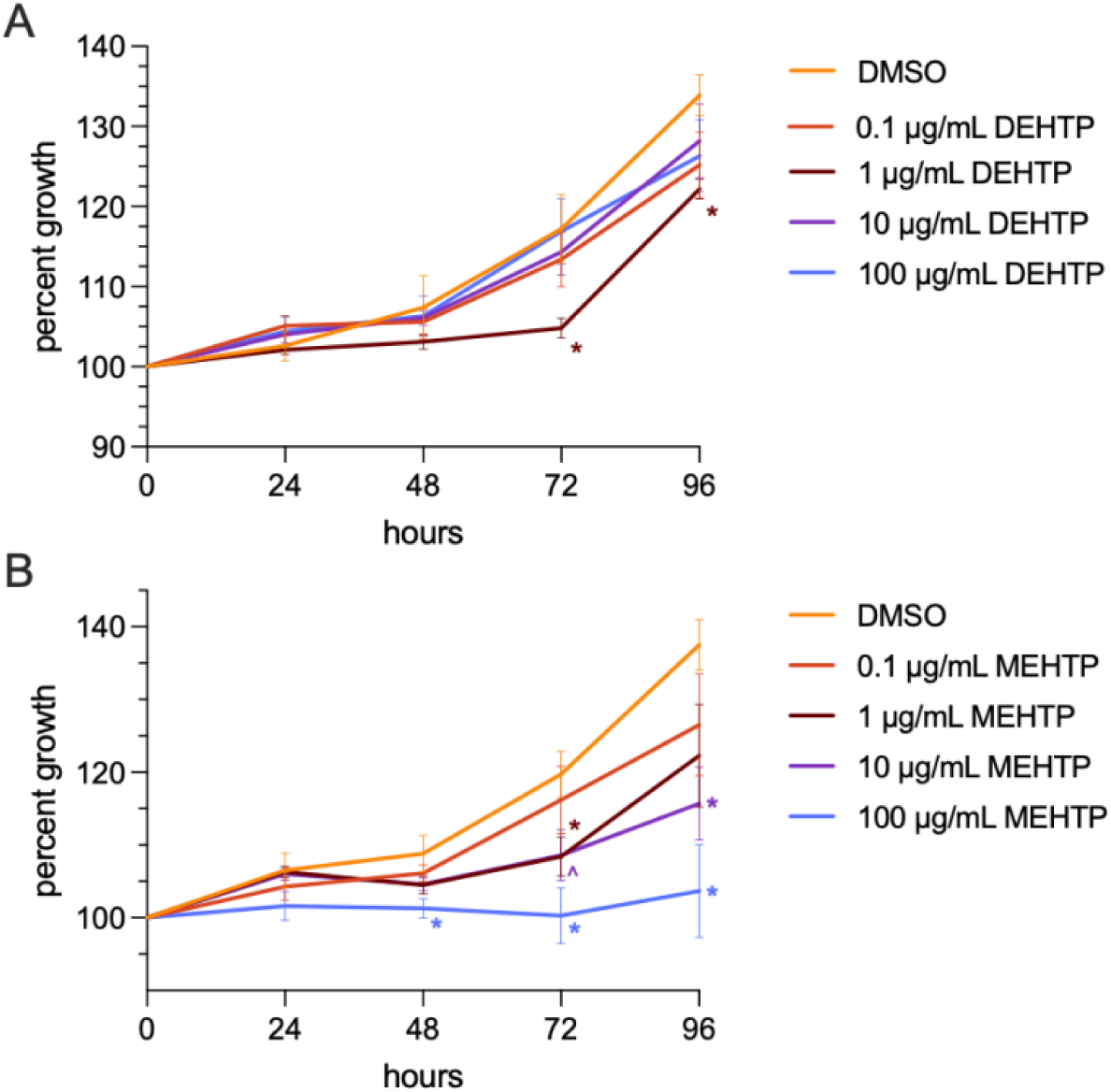
Effects of *in vitro* DEHTP (A) and MEHTP (B) treatment on antral follicle growth. Follicle growth was measured every 24 hrs for 96 hrs. Graphs represent means ± SEM from 5-6 independent experiments per treatment group. Asterisks (∗) indicate significant differences from the control (p ≤ 0.05) and ^ indicates a trend toward significance (p ≤ 0.10).

### 3.6 Effects of DEHTP and MEHTP exposure on gene expression of cell cycle and apoptosis regulators in vitro

DEHTP exposure did not significantly alter expression of cell cycle regulators (**Figure 6A**). Exposure to 10 and 100 µg/ml of MEHP significantly upregulated the cell cycle inhibitor *Cdkn2b* compared to control (**Figure 6B**). DEHTP exposure borderline decreased expression of apoptosis regulator *Casp3* at 1 µg/ml compared to control (**Figure 7A**) and MEHTP exposure borderline increased expression of *Casp8* at 1 µg/ml compared to control (**Figure 7B**). Although the pro-apoptotic factor *Bax* and the anti-apoptotic factor *Bcl2* were not significantly impacted, the ratio of *Bax/Bcl2* expression increased in favor of apoptosis at the 10 and 100 µg/ml MEHTP treatment groups compared to control.

**Figure 6:**
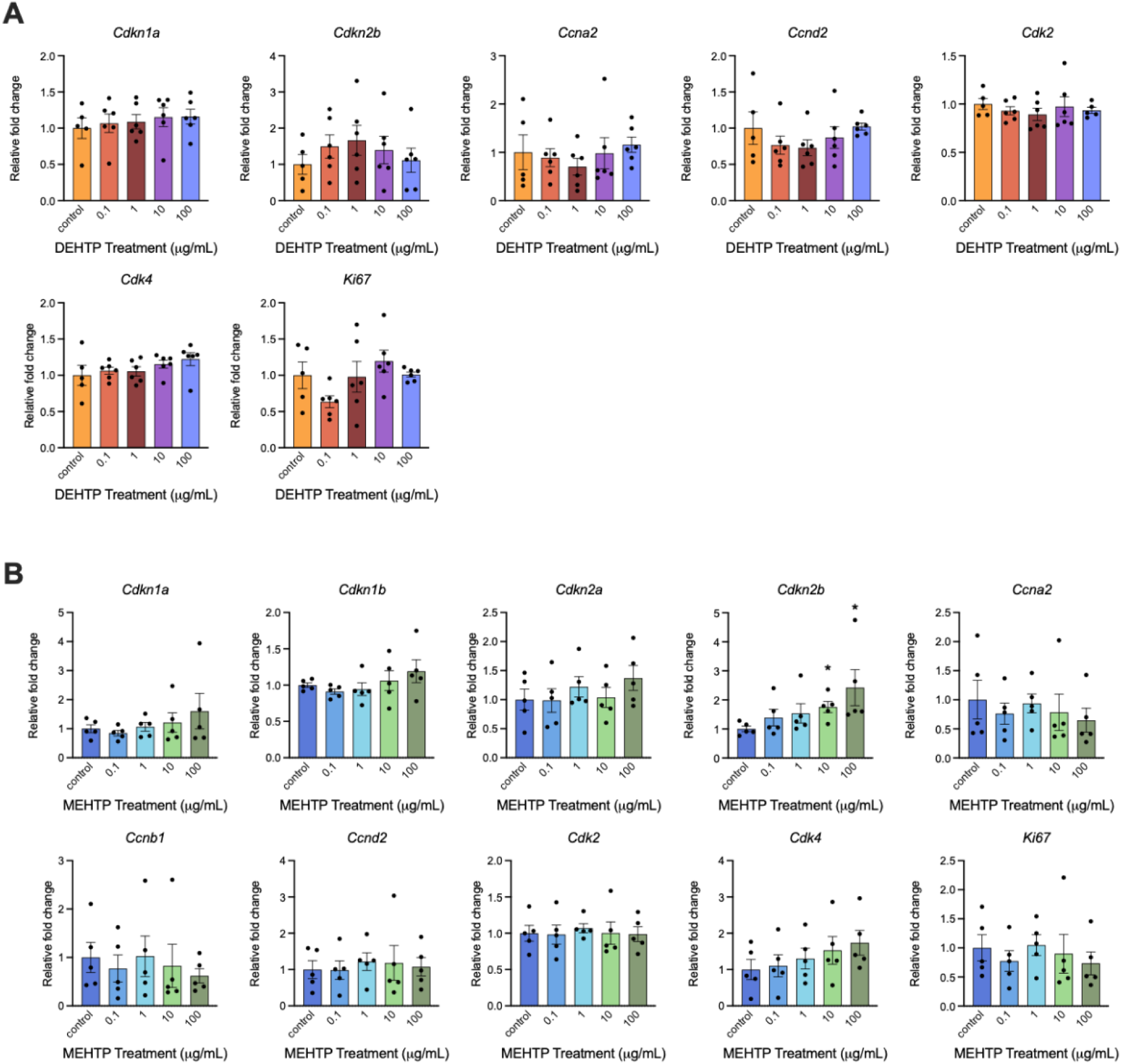
Effects of *in vitro* DEHTP (A) and MEHTP (B) exposure on cell cycle regulator gene expression in antral follicles expressed as fold change compared to vehicle control. All gene expression is relative to the housekeeping gene *BAct*. Graphs represent means ± SD from 5-6 independent experiments per treatment group. Asterisks (∗) indicate significant differences from the control (p ≤ 0.05) and ^ indicates a trend toward significance (p ≤ 0.10).

**Figure 7:**
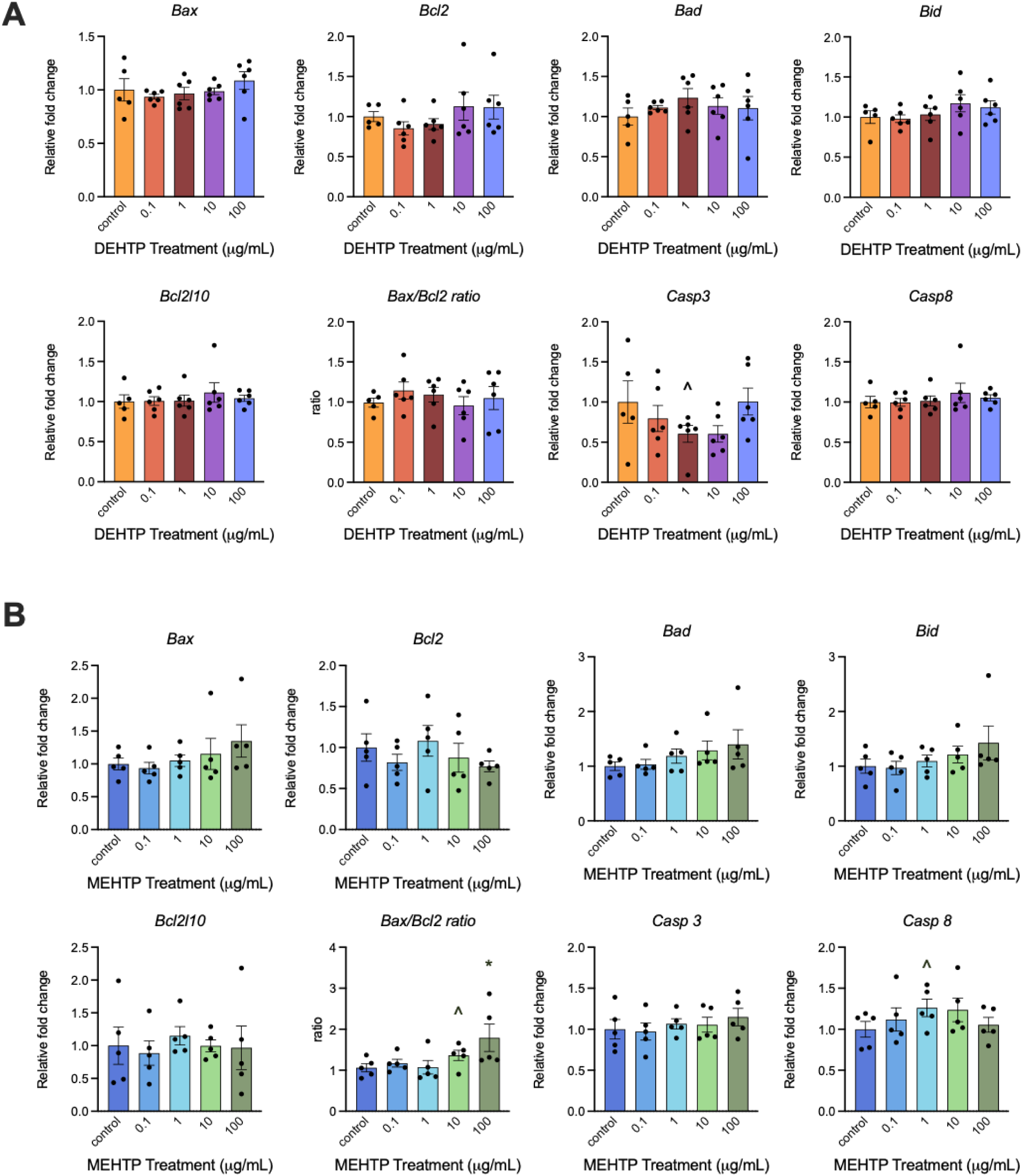
Effects of *in vitro* DEHTP (A) and MEHTP (B) exposure on apoptosis gene expression in antral follicles expressed as fold change compared to vehicle control. All gene expression is relative to the housekeeping gene *BAct*. Graphs represent means ± EM from 5-6 independent experiments per treatment group. Asterisks (∗) indicate significant differences from the control (p ≤ 0.05) and ^ indicates a trend toward significance (p ≤ 0.10).

### 3.7 Effects of DEHTP and MEHTP on sex steroid hormone levels and steroidogenic enzyme gene expression in vitro

DEHTP exposure for 96 hrs did not statistically significantly alter levels of progesterone, androstenedione, testosterone, or estradiol (**Figure 8A**). MEHTP exposure of 100 µg/mL for 96 hrs statistically significantly increased progesterone levels and borderline significantly decreased estradiol levels compared to control (**Figure 8B**). MEHTP exposure for 24 hours did not alter hormone levels (**Figure 8C**).

**Figure 8:**
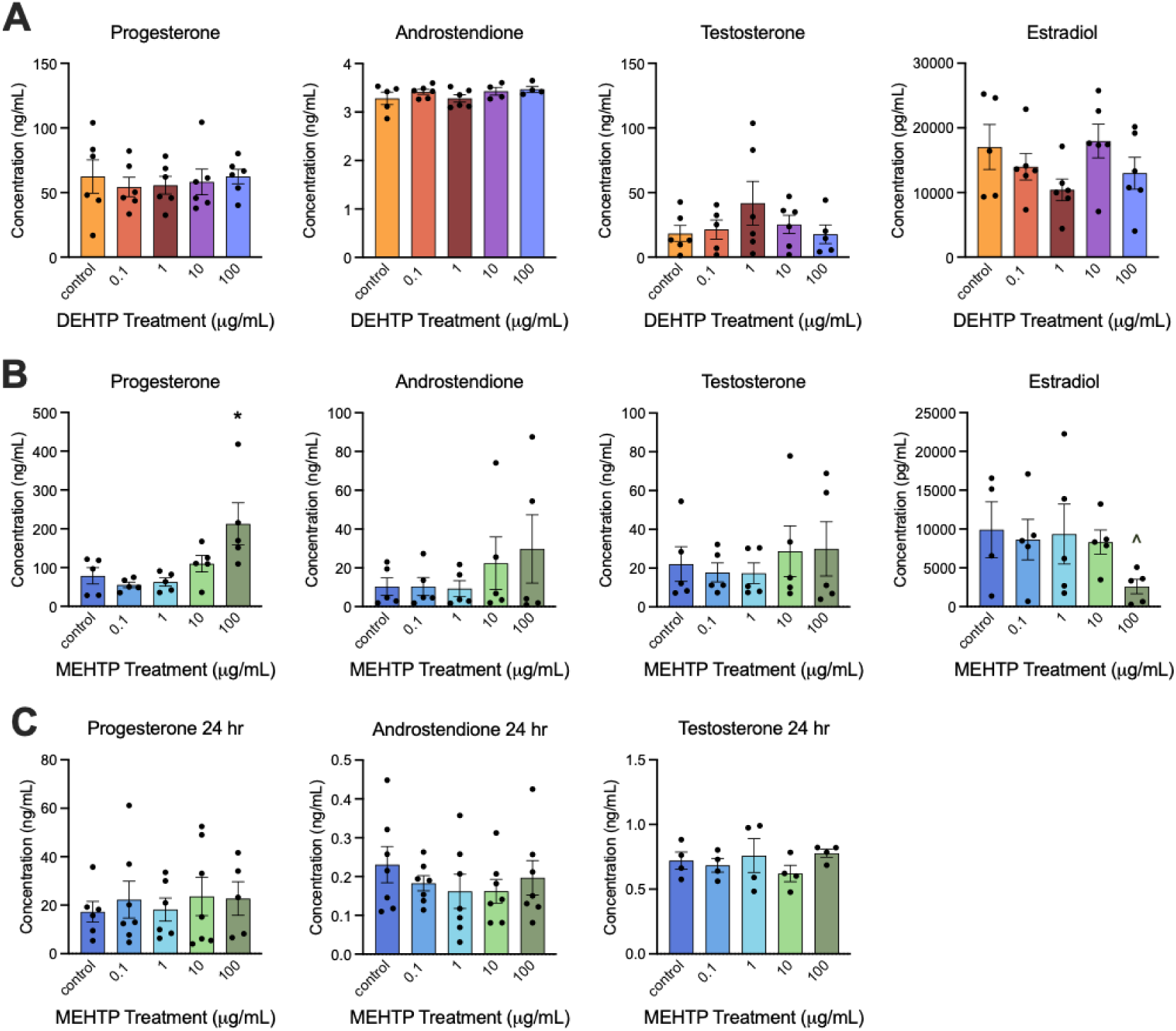
Effects of *in vitro* DEHTP (A) and MEHTP (B) treatment for 96 hours or MEHTP treatment for 24 hours (C) on sex steroid hormone level production by antral follicles. Culture media was subjected to enzyme-linked immunosorbent assays. Estradiol concentrations were too low to be measured after 24 hours. Graphs represent means ± SEM from 4-6 independent experiments per treatment group. Asterisks (∗) indicate significant differences from the control (p ≤ 0.05) and ^ indicates a trend toward significance (p ≤ 0.10).

Exposure to DEHTP for 96 hrs did not alter steroidogenic enzyme gene expression (**Figure 9A**). MEHTP exposure of 10 µg/mL for 96 hrs borderline statistically significantly increased *Cyp11a1* expression compared to control. MEHTP exposure of 10 µg/mL for 24 hrs statistically significantly increased *Star* expression compared to control.

**Figure 9:**
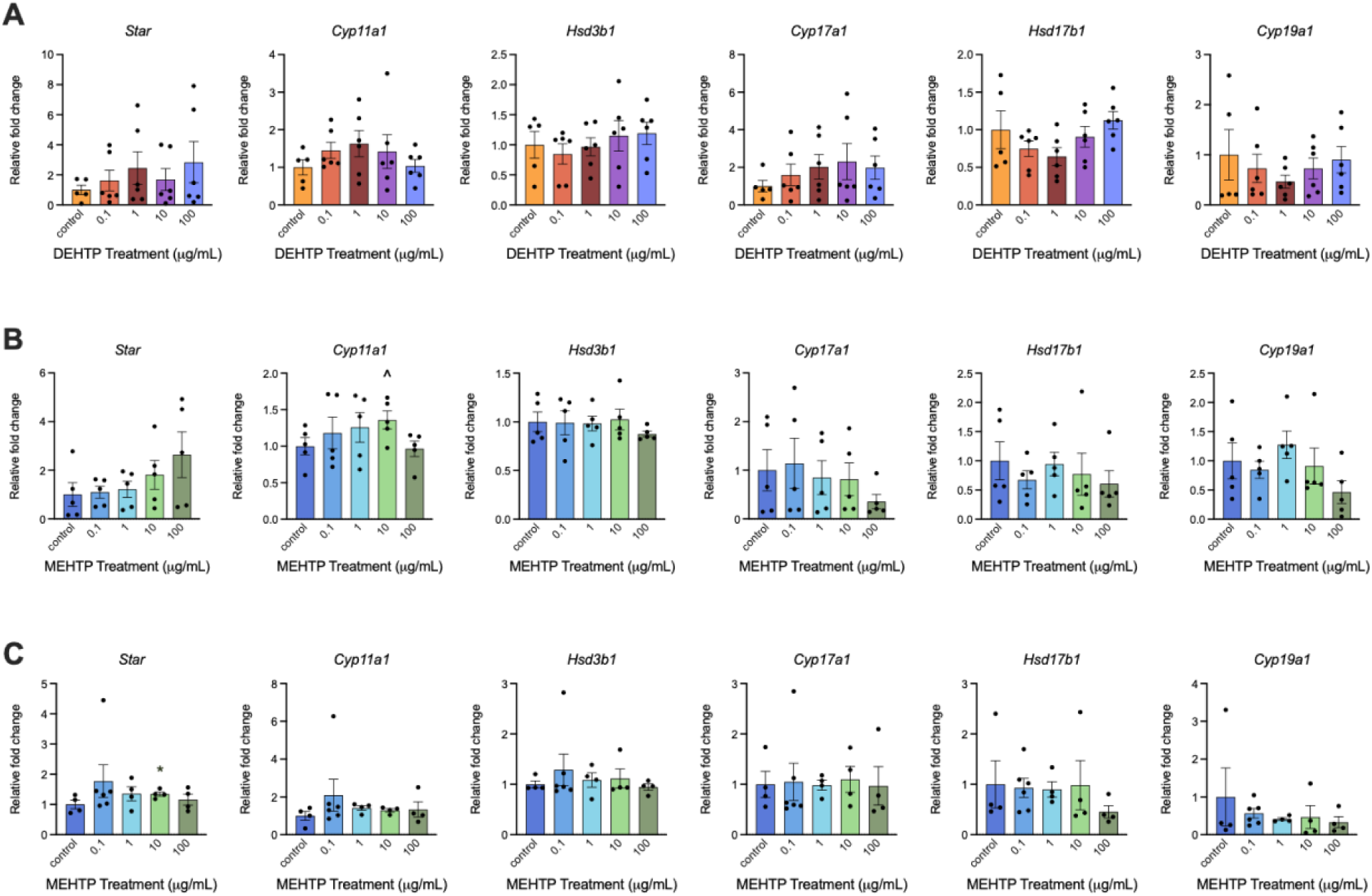
Effects of *in vitro* DEHTP (A) and MEHTP (B) exposure for 96 hrs or MEHTP exposure for 24 hours (C) on steroidogenic enzyme gene expression in antral follicles expressed as fold change compared to vehicle control. All gene expression is relative to the housekeeping gene *BAct*. Graphs represent means ± SEM from 5-6 independent experiments per treatment group. Asterisks (∗) indicate significant differences from the control (p ≤ 0.05) and ^ indicates a trend toward significance (p ≤ 0.10).

## 4. Discussion

This study was performed in response to the pressing need for low dose reproductive toxicological assessment of phthalate replacements with widespread consumer exposure [36]. Many previous studies have shown that phthalates, specifically DEHP and its metabolite MEHP, are ovarian toxicants at low doses [3,37,38]. The phthalate replacement investigated in this study, DEHTP, is a structural isomer of DEHP. Thus, we tested the hypothesis that DEHTP (and its metabolite MEHTP) would have similar ovarian toxicity as DEHP (and its metabolite MEHP). Our results indicated that some levels of DEHTP and MEHTP exposure may disrupt folliculogenesis and steroidogenesis in mouse ovaries. This is important because disruption of normal ovarian function can impair fertility and cause premature reproductive senescence. To our knowledge, this is the first study to investigate the reproductive toxicity of DEHTP at human-relevant exposure levels using a mammalian model.

In this study, we used both *in vivo* and *in vitro* models. For the *in vivo* experiments, we dosed mice with DEHTP directly, whereas for the *in vitro* studies we treated with both DEHTP and its metabolite MEHTP. Phthalates are easily metabolized in the body to monoesters, which are active toxic metabolites [3,39,40]. Toxicokinetic and biomonitoring studies show that DEHTP is biotransformed into analogous para-substituted isomers of DEHP metabolites [6,28]. As both diesters and monoester likely reach the ovary, with varying levels of biotransformation happening on the way or at the ovary based on different routes of exposure, these are both environmentally relevant exposures [41].

Exposure to DEHTP altered follicle counts in ovaries of exposed animals compared to unexposed controls, suggesting that DEHTP exposure disrupts folliculogenesis. The percent of primary follicles was significantly increased in both the 100 µg/kg and 100 mg/kg treatment groups. The corresponding decrease in primordial follicles for the 100 mg/kg treatment group suggests that recruitment of primordial follicles to primary follicles may be accelerated. However, the acceleration of folliculogenesis does not appear to extend beyond primary follicles, as preantral follicles are borderline decreased in the 100 µg/kg treatment group compared to control. This pattern is similar to previously observed effects of 10 day oral exposure to DEHP, which decreased primordial follicle counts at 20 and 200 mg/kg and increased primary follicles at 20 and 200 µg/kg and 20 and 200 mg/kg compared to control [29]. We also observed an increase in the number of abnormal follicles, which were characterized by shrunken oocytes that stain darkly with eosin (pink), at 100 mg/kg. The increase in abnormal follicles rather than preantral follicles suggests that accelerated follicles are not progressing through folliculogenesis but are instead damaged. As the follicle population cannot be renewed, DEHTP exposure may be irreversibly depleting the pool of follicles, which could contribute to infertility and premature ovarian failure. Indeed, when mice dosed with DEHP for 10 days were followed into later life, follicle counts, estrous cyclicity, and fertility were impaired compared to untreated controls [42,43]. Similar long-term studies should be performed following DEHTP exposure for comparison.

Phthalates have been shown to disrupt the cell cycle in the ovary, as well as apoptosis and the process of atresia, or programmed follicle death, by pausing the cell cycle [32,33,44]. Impaired progression of the cell cycle may impede folliculogenesis and atresia; cells that are suspended mid cell-cycle can neither grow nor die [32]. The increase in abnormal follicles observed histologically may be due to their inability to complete the apoptotic process. Gene expression analysis of apoptotic and cell cycle factors supports this hypothesis, as *in vivo* DEHTP exposure significantly decreased the expression of both cell cycle and apoptosis promoters. *In vitro*, follicle growth during the culture period was also slowed, with exposure to increasing concentrations of MEHTP resulting in decreased follicle growth. The stratification of growth by treatment mirrors the effects of MEHP in the same system [37,45]. Interestingly, DEHTP exposure only slowed the growth of follicles exposed to the middle dose. Gene expression analysis of cell cycle and apoptosis regulators *in vitro* did not reveal many statistically significant changes. Of note, the expression of cell cycle inhibitor *Cdkn2b* was upregulated at the 10 and 100 µg/mL doses. Inhibition of the G1/S transition by increased inhibitor expression has previously been observed in phthalates [32,33,44]. Steroid hormone production is an important ovarian function and if disrupted can impact reproductive and non-reproductive health [34]. 10 days of DEHTP exposure resulted in a trend toward decreased testosterone levels. *In vitro*, MEHTP exposure increased progesterone and decreased estradiol. This is a similar trend to a previous study of DEHP, in which 10 days of 200 mg/kg DEHP increased progesterone levels and decreased estradiol [42]. *In vitro*, DEHP and MEHP alone have been shown to decrease estradiol levels [37], and mixtures of phthalates including DEHP and MEHP have been shown to increase progesterone and testosterone and decrease estradiol [32,44]. These effects have been observed after as little as 24 hours of culture [44], but we did not detect changes in hormone levels after 24 hours of MEHTP culture. We also assessed gene expression of steroidogenesis enzymes and identified trending decreases in *Cyp11a1*, which converts cholesterol to pregnenolone, and *Cyp17a1*, which converts pregnenolone to dehydroepiandrosterone and progesterone to androstenedione, *in vivo*, and *Cyp11a1 in vitro*. Unfortunately, our sample size was small, and we did not have enough serum to evaluate more hormones. Overall, these steroidogenesis data suggest that DEHTP trends toward disruption of steroidogenesis similar to phthalates, although more studies with higher sample size are necessary to confirm this observation.

A noticeable finding of this study is that *in vivo* DEHTP exposure more significantly impacted ovarian function compared to *in vitro* exposure to DEHTP or MEHTP. The reduced sensitivity of *in vitro* follicle culture compared to *in vivo* oral dosing followed by whole ovary evaluation (when both experiment types used environmentally relevant doses) has been observed before [46]. *In vivo* experiments capture the effects on the hypothalamus-pituitary-ovary (HPO) axis as well as stroma and other follicle types present in the ovary. A recent zebrafish experiment found that DEHTP exposure disrupted estradiol levels as well as genes expressed in the hypothalamus and pituitary gland, indicating an unbalanced HPO feedback loop [26]. Future studies should examine the effects of DEHTP on neuroendocrine endpoints in rodents. Another possible explanation for the mismatch between *in vitro* and *in vivo* findings is that MEHTP may not be the most active metabolite of DEHTP. A cell line steroidogenesis assay identified mono-2- ethyl-5-hydroxyhexyl terephthalate (MEHHTP) as the most active DEHTP metabolite in disrupting steroidogenesis [25]. DEHTP and/or MEHTP may also be acting through different mechanisms than phthalates.

In conclusion, this study found that *in vivo* and *in vitro* exposure of DEHTP and/or its metabolite MEHTP disrupted folliculogenesis and steroidogenesis in the mouse ovary. Some, but not all, of the observed results are consistent with previous studies of DEHP and its metabolite MEHP, which are ring substitution isomers of DEHTP and MEHTP. Disruption of ovarian steroidogenesis and folliculogenesis could be contributing to observed epidemiological associations between biomarkers of DEHTP exposure and altered hormone signaling in humans. As the substitution of DEHP and other phthalates with replacements of similar structures continues to increase in consumer products, it is important to evaluate potential health risk of replacements at low doses.

## Data Availability Statement

Data will be made available by the corresponding author upon reasonable request.

## Acknowledgements

Thank you to all members of the EDC Lab at NJIT and the Flaws Lab at the University of Illinois at Urbana-Champaign.

## Author Contributions: Credit

**Courtney Potts**: investigation, project administration, validation, visualization, writing - original draft, writing - reviewing and editing; **Allison Harbolic**: investigation, visualization, writing - original draft; **Maire Murphy**: investigation; **Michelle Jojy**: investigation, validation, visualization; **Christine Hanna**: investigation, validation; **Maira Nadeem**: investigation, validation; **Hanin Alahmadi**: investigation, validation; **Stephanie Martinez**: investigation, validation, writing - reviewing and editing; **Genoa Warner**: conceptualization, methodology, validation, formal analysis, investigation, data curation, writing - original draft, writing - review and editing, visualization, supervision, project administration, funding acquisition

## Abbreviations

DEHTP: Di-2-ethylhexyl terephthalate
MEHTP: Mono-2-ethylhexyl terephthalate
DEHP: di-2-ethylhexyl phthalate
MEHP: mono-2-ethylhexyl phthalate
MEHHTP: mono-2-ethyl-5-hydroxyhexyl terephthalate
DMSO: dimethyl sulfoxide
NHANES: National Health and Nutrition Examination Survey
ELISA: enzyme linked immunosorbent assay

