## supplementary materials for "A common phthalate replacement disrupts ovarian function in young adult mice"

**Table S1:** DNA oligonucleotide primers used for qPCR gene expression assays.

| Gene symbol | Gene name | Forward primer (5'→3') | Reverse primer (5'→3') |
|---|---|---|---|
| *ActB* | Beta actin | GTGACGTTGACATCCGTAAAGA | GCCGGACTCATCGTACTCC |
| *Bad* | BCL2-associated agonist of cell death | TGAGCCGAGTGAGCAGGAA | GCCTCCATGATGACTGTTGGT |
| *Bax* | BCL2-associated X protein | AGACAGGGGCCTTTTTGCTAC | AATTCGCCGGAGACACTCG |
| *Bcl2* | B cell leukemia/lymphoma 2 | GCTACCGTCGTGACTTCGC | CCCCACCGAACTCAAAGAAGG |
| *Bcl2l10* | BCL2-like 10 | CCACTGCATGAACGCACTAGA | GAGAGCAACTTATCTGCCATCTG |
| *Bid* | BH3 interacting domain death agonist | GCCGAGCACATCACAGACC | TGGCAATGTTGTGGATGATTTCT |
| *Casp3* | caspase 3 | CTCGCTCTGGTACGGATGTG | TCCCATAAATGACCCCTTCATCA |
| *Casp8* | caspase 8 | TGCTTGGACTACATCCCACAC | GTTGCAGTCTAGGAAGTTGACC |
| *Ccna2* | cyclin A2 | AAGAGAATGTCAACCCCGAAAAA | ACCCGTCGAGTCTTGAGCTT |
| *Ccnd1* | cyclin D1 | GCGTGTGCCTGTGACAGTTA | CCTAGCGTTTTTGCTTCCCTT |
| *Ccnd2* | cyclin D2 | GAGTGGGAACTGGTAGTGTTG | CGCACAGAGCGATGAAGGT |
| *Cdk2* | cyclin dependent kinase 2 | ATGGAGAACTTCCAAAAGGTGG | CAGTCTCAGTGTCGAGCCG |
| *Cdk4* | cyclin dependent kinase 4 | ATGGCTGCCACTCGATATGAA | TGCTCCTCCATTAGGAACTCTC |
| *Cdkn1a* | cyclin dependent kinase inhibitor 1A | CCTGGTGATGTCCGACCTG | CCATGAGCGCATCGCAATC |
| *Cdkn1b* | cyclin dependent kinase inhibitor 1B | TCAAACGTGAGAGTGTCTAACG | CCGGGCCGAAGAGATTTCTG |
| *Cdkn2a* | cyclin dependent kinase inhibitor 2A | CGCAGGTTCTTGGTCACTGT | TGTTCACGAAAGCCAGAGCG |
| *Cdkn2b* | cyclin dependent kinase inhibitor 2B | CCCTGCCACCCTTACCAGA | GCAGATACCTCGCAATGTCAC |
| *Cyp11a1* | Cytochrome P450, family 11, subfamily a, polypeptide 1 Cytochrome | AGGTCCTTCAATGAGATCCCTT | TCCCTGTAAATGGGGCCATAC |
| *Cyp17a1* | Cytochrome P450, family 17, subfamily a, polypeptide 1 Cytochrome | AGTCAAAGACACCTAATGCCAAG | ACGTCTGGGGAGAAACGGT |
| *Cyp19a1* | Cytochrome P450, family 19, subfamily a, polypeptide 1 Cytochrome | AACCCCATGCAGTATAATGTCAC | AGGACCTGGTATTGAAGACGAG |
| *Hsd17b1* | 17-beta-hydroxysteroid dehydrogenase 1 | TGGCTGTTCGCCTAGCTTC | GGCCTCCAACAATGGTCCC |
| *Hsd3b1* | 3-beta-hydroxysteroid dehydrogenase 1 | AGCTCTGGACAAAGTATTCCGA | GCCTCCAATAGGTTCTGGGT |
| *Ki67* | antigen identified by monoclonal antibody Ki 67 | ATCATTGACCGCTCCTTTAGGT | GCTCGCCTTGATGGTTCCT |
| *Rn18s* | 18S ribosomal RNA | TCGGCGTCCCCCAACTTCTTA | GGTAGTAGCG ACGGGCGGTGT |
| *Star* | Steroidogenic acute regulatory protein | CGGGTGGATGGGTCAAGTTC | GCACTTCGTCCCCGTTCTC |

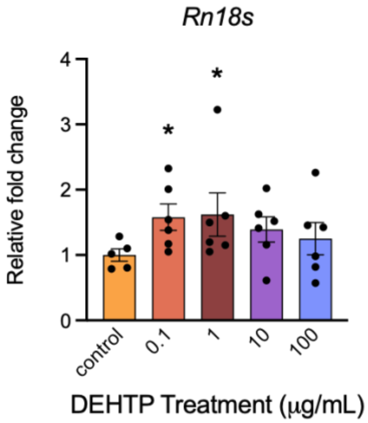


**Figure S1**: Effects of *in vitro* DEHTP exposure on *Rn18s* gene expression in antral follicles expressed as fold change compared to vehicle control. All gene expression is relative to the housekeeping gene *BAct*. Graph represents means ± SEM from 5-6 independent experiments per treatment group. Asterisks (∗) indicate significant differences from the control ($p \leq 0.05$).

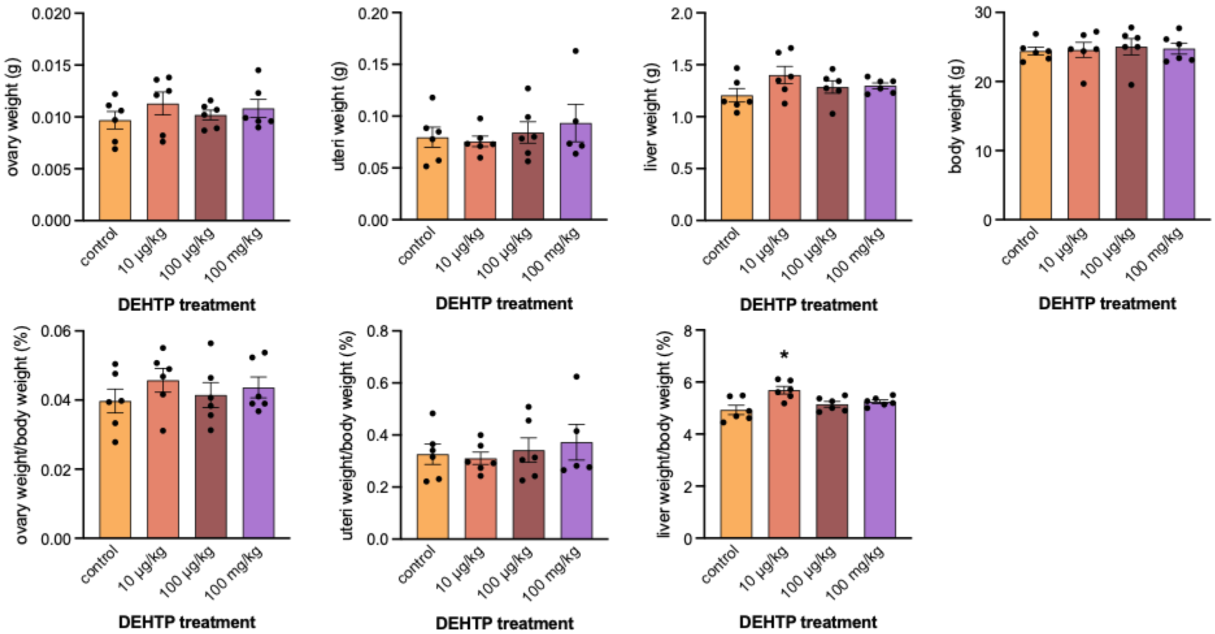


**Figure S2**: Effects of adult exposure to DEHTP on ovary weight, uteri weight, liver weight, body weight, and the percent weight of each organ normalized to body weight in female mice. Graphs represent means ± SEM from 5–6 animals per treatment group. Asterisk (*) indicates significant difference from the control ($p \leq 0.05$).
